# Prenatal assembly of functional cortical circuits

**DOI:** 10.64898/2026.06.01.729224

**Authors:** Ilaria Morassut, Alessandra Panzeri, Sabine Fièvre, Julien Prados, Samir Vaid, Florent Montange, Adrien Debry, Jiafeng Zhou, Riccardo Bocchi, Esther Klingler, Camilla Bellone, Athanasia C. Tzika, Denis Jabaudon, Sergi Roig-Puiggros

## Abstract

The extent to which early brain maturation requires interactions with the outside world is a central question in neurobiology. Because much of cortical maturation occurs after birth in *Mus*, it has often been viewed as dependent on postnatal experience. However, studies addressing this issue have largely relied on postnatal sensory deprivation paradigms, which perturb normal development and cannot determine to what extent postnatal experience *per se* drives maturation. To address this question, here we compare two related rodents with markedly different gestation lengths: the precocial *Acomys dimidiatus* (39-day gestation) and the altricial *Mus musculus* (19-day gestation). By generating a novel *Acomys* reference genome and using histology, cellular birth dating, electrophysiology, single-nucleus transcriptomics, and quantitative behavior, we show that *Acomys* preserves the canonical sequence and timing of cortical development, while shifting major milestones of neuronal, circuit and behavioral maturation into prenatal life. At birth, *Acomys* cortex already shows advanced cytoarchitecture, neuronal physiology, thalamocortical barrels, transcriptional states, and sensorimotor behavior, with postnatal molecular programs in *Mus* unfolding prenatally in *Acomys*. Thus, birth is not a prerequisite for early cortical maturation. Instead, evolutionarily conserved developmental programs unfold across birth, with birth occurring at different stages of these programs, reflecting a species-specific balance between neonatal competence and prolonged postnatal plasticity.

## Introduction

A fundamental question in developmental neuroscience is how cortical circuits acquire functional maturity. In altricial mammals, including humans and the laboratory mouse (*Mus musculus*, hereafter referred to as *Mus*), the cerebral cortex is immature at birth and undergoes extensive postnatal refinement^1^. During this period, sensory-driven activity shapes synaptic connectivity^2–4^, guides the stabilization and elimination of synapses^5^, and defines critical periods of heightened plasticity^6^. Landmark studies in visual and somatosensory systems, mostly based on lesion of sensory organs have established experience-dependent mechanisms as central to cortical development^7–9^. These studies, together with the temporal coincidence of birth and the onset of neuronal maturation in altricial species, have established the view that cortical maturation programs rely on postnatal interactions. However, in precocial species, newborns exhibit advanced motor coordination and sensory capabilities at birth, suggesting that substantial neural maturation can occur prenatally^10,11^. This raises the possibility that cortical developmental programs are broadly conserved across species, with birth occurring at different stages along a shared maturation trajectory. Testing this idea requires comparing related species with aligned cortical cell types but different gestation lengths. Precocial mammals thus provide an opportunity to disentangle the relative contributions of intrinsic developmental programs and postnatal experience to cortical maturation and function.

Towards this goal, here we investigate the precocial rodent *Acomys dimidiatus* (hereafter referred as *Acomys*), characterized by extended gestation and advanced maturity at birth. As a phylogenetically close relative of *Mus* (adjusted divergence time 12.4M years^12^, analogous to that separating *Mus* and *Rattus*), *Acomys* provides a comparative model to determine the extent to which cortical maturation programs require postnatal experience, or can unfold prenatally when birth occurs later relative to development.

We combine histological, single-nucleus transcriptomic, electrophysiological, and behavioral approaches to compare *Mus* and *Acomys* development and determine how prenatal and postnatal environments shape cortical circuit assembly and maturation. We find that alongside advanced sensorimotor function present compared to *Mus*, prolonged prenatal development in *Acomys* follows canonical *Mus* timescales *i*.*e*. the neurogenic period is 7 days-long, neurons follow conserved transcriptional and morphological maturation trajectories, and thalamocortical circuits assemble into barrels. Precocial development therefore reflects a prenatal shift of conserved maturation programs, suggesting that key aspects of cortical function can emerge before postnatal sensory experience.

## Results

### *Acomys* displays hallmarks of precocial maturity at birth

*Mus* newborns are altricial. They lack fur, have closed eyes and exhibit limited motor abilities. Despite whiskers being present before birth^13^, they mostly elongate during the first postnatal week^14^, coinciding with the maturation of the whisker-to-barrel pathway^15^. In *Mus*, this early postnatal period represents a developmental window during which whisker-related sensory experience and activity-dependent mechanisms are known to shape the structural and functional maturation of barrel-cortex circuits^16^. Indeed, active whisking behavior, for detecting objects in space^17^, only emerges from postnatal day (P) 12 onwards^18^, reflecting both circuit maturation and the onset of exploratory behavior.

In contrast, *Acomys* newborns are precocial. They are fully furred, have open eyes, are mobile, and possess elongated whiskers at birth^19,20^ (**Fig. 1a**). When presented with an object, *Acomys* newborns actively whisk to explore it as early as P1 (**Fig. 1a, Supplementary information video 1**), suggesting that they possess sensorimotor capabilities corresponding to late postnatal stages in *Mus* (**Supplementary information video 2**). This maturity at birth prompted us to ask whether developmental events unfolding postnatally in *Mus* are shifted into prenatal life in *Acomys*.

**Figure 1.**
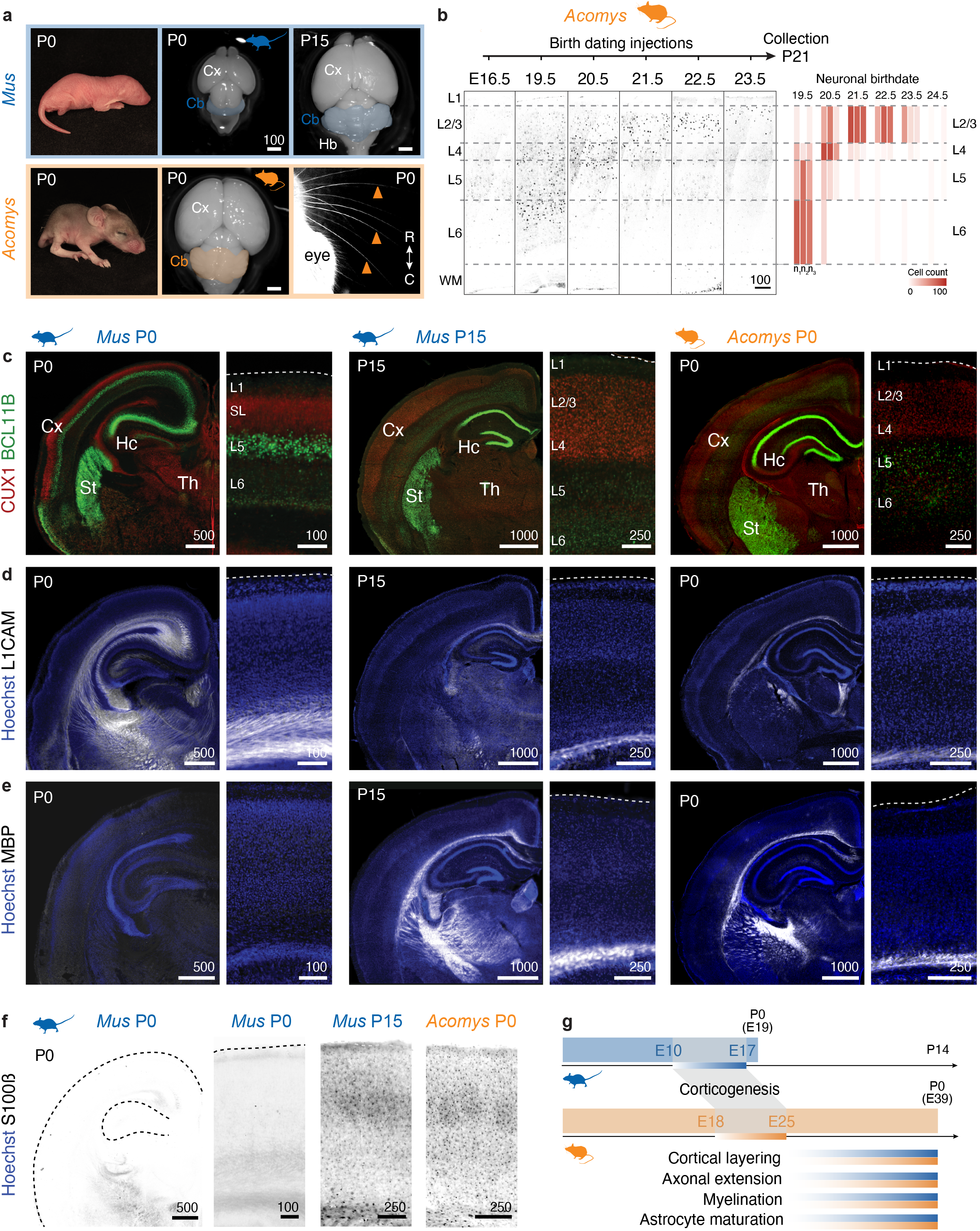
The brain of the precocial rodent *Acomys dimidiatus* exhibits mature-like features at birth. **a**, Left, images of *Mus* (top) and *Acomys* (bottom) newborns at PO. Right, images of PO and P15 *Mus* brain (top) and *Acomys* PO brain and whiskers (bottom). The blue shadow in the *Mus* images and the orange shadow in the *Acomys* image highlight the cerebellum (Cb). Arrowheads indicate individual whiskers in *Acomys* pups at PO. **b**, Birth dating timeline of *Acomys dimidiatus*. Left, immunofluorescence images of P21 brains injected with thymidine analogs at the indicated timepoints. Right, percentage of cells positive for thymidine-analog labeling in each cortical layer across timepoints and replicates (columns). **c-f**, CUX1 and BCL11B, (**c**), L1CAM (**d**), MBP (**e**), S100B (**f**) immunofluorescence for *Mus* coronal sections at PO and P15, and *Acomys* at PO. Scale bars measure unit: µm. **g**, Schematic of developmental timelines for *Mus* (top, in blue) and Aco*mys* (bottom, in orange) indicating main corticogenesis and maturation events in the respective species. Gradients of orange and blue represent progressive emergence of the respective feature. Abbreviations: Cx, cortex; Cb, cerebellum; E, embryonic day; P, postnatal day; Th, thalamus.

### Extended gestation in *Acomys* does not extend cortical neurogenesis

*Acomys* has a gestational length of approximately 39 days^19^ compared to the 19 days in *Mus*. Neurogenesis has been shown to start later in precocial species including *Acomys cahirinus*^10^, for which corticogenesis has been shown to start at E18^21^. To determine the onset and duration of corticogenesis in *Acomys*, we performed birth dating experiments using thymidine analogs in pregnant dams from embryonic day (E) 16.5 to E24.5 (**Methods**) and analyzed offspring cortices by immunofluorescence at postnatal day P21 (**Fig. 1b**). Surprisingly, although gestation is almost twice as long in *Acomys* compared to *Mus*, corticogenesis lasts only 7 days, which is similar to what happens in *Mus*^22^, with onset around E18^21^, ending by E24.5, and following an inside-out pattern of neuronal generation (**Fig. 1b**). Thus, by the end of neurogenesis, *Acomys* embryos have a 14-day window for prenatal neuronal maturation, a process that occurs postnatally in *Mus*. Based on this finding, we established a comparative developmental timeline aligning *Acomys* E24.5 with *Mus* E17 (*i*.*e*., end of corticogenesis in both species), allowing systematic comparison of cortical maturation before vs. after birth for equivalent developmental features.

### *Acomys* brain matures extensively before birth

At birth, the *Acomys* brain weighs approximately 55% of its adult weight, compared to only about 20% for *Mus*^19^. Consistent with an advanced developmental state, the *Acomys* cerebellum is already foliated at P0, a feature that in *Mus* typically emerges during postnatal development and is evident by P15 (**Fig. 1a**). To determine whether cortical development displays a similar degree of precocity, we examined cortical layering, axonal extension, myelination and astrocytic colonization in *Acomys* (**Fig. 1c-f**).

Completion of neurogenesis and neuronal migration establish distinct cortical laminae characterized by the expression of specific molecular markers^23^. At P0, *Mus* cortical layers are still poorly defined due to ongoing neuronal migration but are clearly visible at P15^24^. To assess *Acomys* lamination at birth, we performed immunostainings against BCL11B (expressed by layer (L)5b cortical neurons)^25,26^ and CUX1 (expressed by superficial layer neurons)^27,28^. At birth, *Acomys* cortices exhibited a six-layered structure with mutually exclusive and segregated CUX1/BCL11B expression pattern, comparable to *Mus* at P15 (**Fig. 1c**).

During *Mus* early postnatal development, axons from thalamocortical and cortical projecting neurons still extend towards their targets (refs). L1CAM, a neuronal cell adhesion molecule crucial for axon growth, fasciculation, and fiber tract formation during this period^29,30^, is highly expressed by *Mus* P0 axons (**Fig. 1d** left), which in turn, are not yet myelinated by oligodendrocytes^31^ (myelin basic protein (MBP), **Fig. 1e** left). By P15, L1CAM is confined to MBP-positive axon bundles, such as the corpus callosum and the internal capsule, consistent with the completion of large-scale axonal growth and pathway stabilization (**Fig. 1d-e** middle). At birth, *Acomys* displayed L1CAM and MBP patterns resembling that of P15 *Mus*, with organized, well-defined, and myelinated axonal tracts, suggesting that major axonal projections are already established at birth in this species (**Fig. 1d-e** right). In *Mus* cortex, astrocytes are sparse at birth, as astrogenesis has just begun and clonal expansion within the cortical parenchyma occurs over the first two postnatal weeks^33^. Accordingly, S100β-positive astrocytes form a dense and evenly spaced network across cortical layers by P15^32,34,35^ (**Fig. 1f**). In contrast, at P0, *Acomys* already displayed abundant S100β-positive astrocytes throughout the cortex, with a distribution pattern resembling that of *Mus* at P15 (**Fig. 1f** right). Together, these results show that major anatomical hallmarks of cortical maturation are established before birth in *Acomys* (**Fig. 1g)**.

### Cortical neurons acquire postnatal-like morpho-functional properties before birth in *Acomys*

Next, we asked whether this anatomical maturity was accompanied by corresponding morphological and electrophysiological maturation of cortical glutamatergic neurons. For this purpose, we combined whole-cell patch-clamp recordings with biocytin filling of single L5b extratelencephalically-projecting and L2/3 intratelencephalically-projecting neurons in the developing cortex of *Mus* and *Acomys* (**Fig. 2a, Extended Data Fig. 1a**).

**Figure 2.**
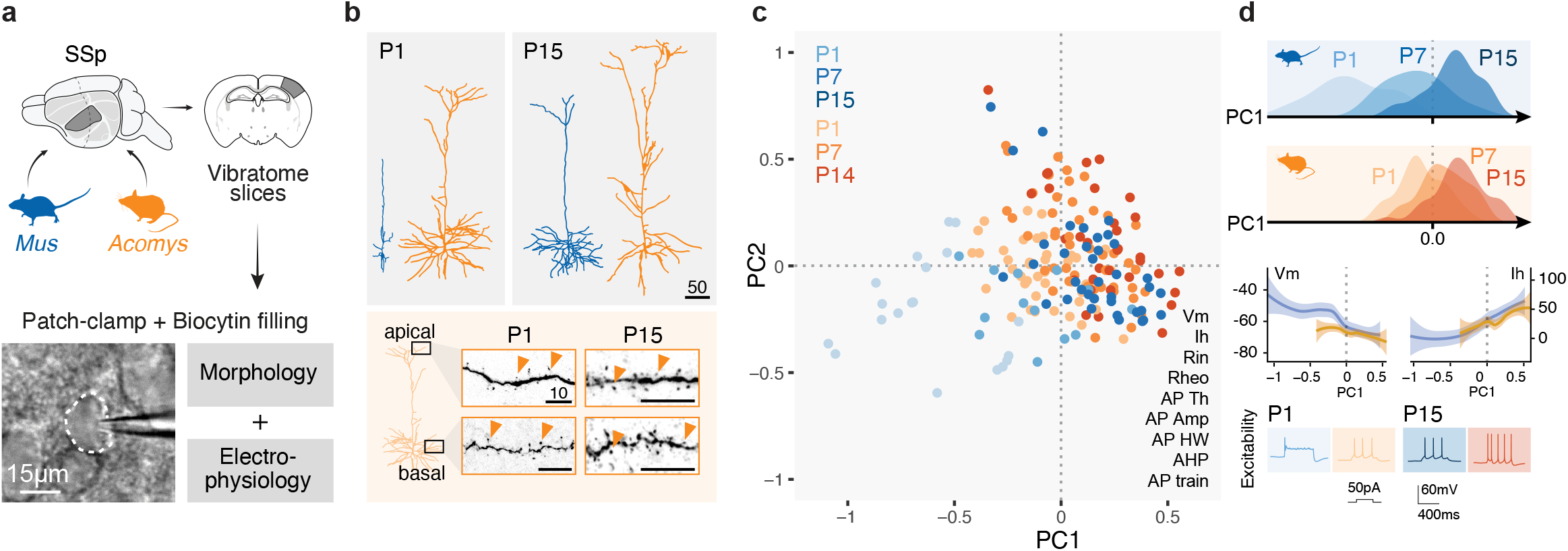
*Acomys* cortical neurons display developmentally advanced morphological and electrophysiological features at birth. **a**, Schematic representation of the experimental strategy. Bottom: photomicrograph of a patched cell. **b**, Top, examples of morphologically reconstructed *Mus* (blue) and *Acomys* (orange) L5 ET glutamatergic neurons at the indicated ages. Bottom, examples of dendritic spines appearance in *Acomys* neurons at the indicated ages. Scale bar measure unit: µm. **c**, Principal Component Analysis (PCA) plot of *Acomys* (orange) and *Mus* (blue) neurons, for the 9 electrophysiological parameters listed on the bottom right. **d**, Top, density plot along PC1 axis for *Mus* neurons (top, in blue) and *Acomys* neurons (below, in orange) at P1, P7 and P15. Middle, values of membrane potential (Vm) and h current (lh) are plotted as examples of dynamically varying parameters along PC1. Bottom, examples of excitability traces for *Mus* neurons (in blue) and *Acomys* neurons (in orange-red) at P1 (left) and P15 (right), showing mature-like patterns in *Acomys* P1 neurons. Abbreviations: SSp, primary somatosensory cortex; PC, principal component; Vm, resting membrane potential; lh, hyperpolarization-activated current; Rin, input resistance; Rheo, rheobase; AP Th, action potential threshold; AP amp, action potential amplitude; AP HW, action potential half-width; AHP, hyperpolarization potential; AP train, action potential train.

At P1, some *Mus* projection neurons are still migrating^36^ and exhibit simple morphologies with short apical dendrites and minimal basal arbors, both in deep layers (**Fig. 2b**) and superficial layers (**Extended Data Fig. 1b**). Over the first two postnatal weeks, they extensively develop their apical and basal dendritic arbors^37^, reaching mature morphologies by P15 in the primary somatosensory (SSp) cortex (**Fig. 2b, Extended Data Fig. 1b)**. In *Acomys*, neurons at E25 displayed immature dendritic morphologies (**Extended Data Fig. 1b, c**). However, at P1, *Acomys* neurons already exhibited a dendritic complexity comparable to *Mus* P15 neurons, with well-elaborated apical and basal arbors in both layer 5b and layer 2/3 (**Fig. 2b, Extended Data Fig. 1b, c**). Furthermore, *Acomys* neuronal dendrites exhibited dendritic spines (**Fig. 2b, Extended Data Fig. 1b, c** bottom insets), indicating the presence of morphological synapses, which only occur after the second postnatal week in *Mus*^38^.

Accordingly, electrophysiological recordings revealed that *Acomys* neurons undergo substantial functional maturation *in utero* (**Supplementary information table 1)**. Using an unsupervised approach (Principal Component Analysis, PCA), *Mus* and *Acomys* neurons were ordered based on the co-variation of nine electrophysiological parameters (**Fig. 2c, Extended Data Fig. 1e**). The unbiased alignment of all the recorded neurons across the first principal component (PC1) reflected their postnatal maturation process (**Fig. 2d** top). At birth, *Mus* neurons presented immature firing behavior, characterized by low excitability and broad, slowly repolarizing action potentials. By P14, *Mus* neurons fired repetitive trains of narrow spikes with higher frequency and amplitude, reflecting progressive increase of membrane excitability and reduced input resistance (**Fig. 2d** middle). In *Acomys*, however, P1 neurons already exhibited mature firing patterns (**Fig. 2d** bottom) and passive membrane properties (**Extended Data Fig. 1f**), consistent with acquisition of mature electrophysiological competence. Based on PC distances, *Acomys* neurons display *Mus* P7-like electrophysiological features already at P1 (**Fig. 2d** top), supporting a substantial level of prenatal acquisition of electrophysiological properties that are acquired postnatally in *Mus*.

### Postnatal *Mus* transcriptional programs unfold prenatally in *Acomys*

To molecularly characterize the effects of prenatal maturation on cortical cell types, we performed single-nuclei RNA sequencing of the *Acomys* SSp cortex at five developmental stages: E25, P0, P7, P14 and adult (P113). For comparison with *Mus*, we used available SSp datasets spanning from E17 to adult^39,40^ (**Fig. 3a**). We generated a high-quality *Acomys dimidiatus* reference genome to enable transcriptomic profiling of *Acomys* at a quality comparable to *Mus* (**Extended Data Fig. 2a**): transcripts were annotated using bulk RNA sequencing data from multiple organs (**Supplementary Information Table 2**) and processed with the “EGAPx” pipeline (**Extended Fig. 2a, Methods**). Following quality control, each *Acomys* and *Mus* timepoint dataset was independently annotated using “MapMyCells”^41^ (Allen Brain Institute). The annotated datasets were then integrated using a SysVI-based model^42^ (**Fig. 3b, Extended Data Fig. 2b-d**).

**Figure 3.**
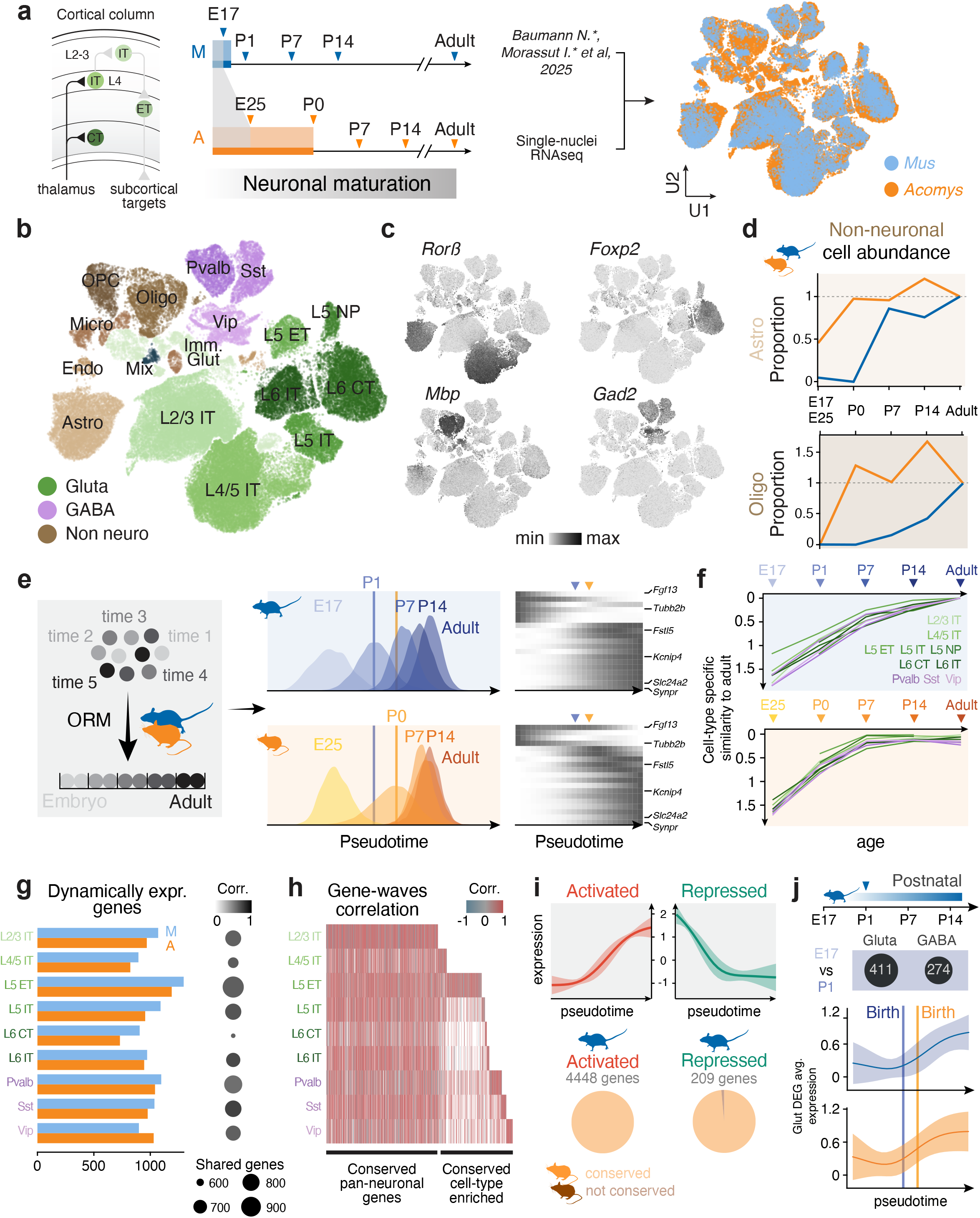
Postnatal *Mus* maturation programs occur prenatally in *Acomys*. **a**, Left: schematics of S1 circuit with involved glutamatergic cell types. Middle: timeline of published datasets used for the analysis for *Mus* (top) and *Acomys* dataset generation (bottom). Right: UMAP of integrated datasets of *Mus* and *Acomys* (color-coded). **b**, UMAP representation of cell type annotation. **c**, Feature plots of key representative marker genes. **d**, Relative proportions of Astrocytes (top) and Oligodendrocytes (bottom), color-coded per species. **e**, Left: schematic representation of the pseudotime modeling used and resulting density plots for *Mus* (top, in blue) and *Acomys* (bottom, in orange). Progressive ages are color-coded with increasing shades. Circles in schematic represent individual cells. Right: heatmap representation of gene waves for corresponding model genes in glutamatergic neurons for *Mus* top, and *Acomys* bottom, with representative gene names indicated on the right. Arrowheads indicate the peak for *Mus* P0 (blue arrowhead) and *Acomys* P1 (orange arrowhead). **f**, Earth’s mover distance measured relative to the *Mus* adult pseudotime distribution for *Mus* (top) and *Acomys* (bottom) cell types (color-coded). **g**, Bar plots of dynamically expressed genes in *Mus* and *Acomys* (color-coded) per cell type (columns). Dot plots indicate the number of shared genes between the two species (size of the dots) and their average cross-species temporal correlation value (gray scale). **h**, Heatmap of correlation values (color-coded) for each cell type (rows) and gene wave (columns). The conserved genes across cell type cluster on the left side of the heatmap. **i**, Top right: average wave profile for each type of dynamics (left, Activated; right, Repressed). Pie charts of proportions of genes with Activated dynamics in *Mus* that are showing the same (conserved) or different (not conserved) dynamics in *Acomys*. **j**, Top, dot plot of DEGs between *Mus* E17 vs P1 in glutamatergic and GABAergic neurons. Average wave profile of DEGs identified in E17 vs P1 comparison in *Mus*, for *Mus* (top) and *Acomys* (bottom) for glutamatergic neurons. Abbreviations: A, Acomys; Astro, astrocytes; Oligo, oligodendrocytes; OPC, oligodendrocyte precursor cells; GABA, GABAergic neurons; Gluta, Glutamatergic; CTX, cortex; lT, intratelencephalically-projecting neurons; ET, extratelencephalically-projecting neurons; M, Mus; ORM, ordinal regression model; P, postnatal day; U, UMAP.

UMAP embedding revealed conserved neocortical cell types between species, including glutamatergic, GABAergic and non-neuronal populations (**Fig. 3a-c, Extended Data Fig. 2e-g**). In *Mus*, glial colonization of the developing cortex occurs postnatally. Fiber tract myelination by oligodendrocytes begins at P5, and astrocyte subtypes expand until P7^33^. Accordingly, cell type proportions shift substantially from P0 to P7 with the emergence of non-neuronal types (**Fig. 3d, Extended Data Fig. 2e-g**). In contrast, *Acomys* P0 samples displayed a cellular composition more closely resembling older *Mus* brains, with a higher proportion of non-neuronal populations (**Fig. 3d, Extended Data Fig. 2e-g**). Fluctuations in relative proportions between successive stages may also partly reflect the greater fragility of mature neurons compared with glial cells during cell dissociation at mature stages. Nevertheless, these data confirming observations made at the histological level (**Fig. 1e, f**). Finally, *Mus* astrocytes undergo molecular maturation from E16 to P21 (**Extended Data Fig. 3a, b**), which we captured as sequential gene expression waves, each containing temporally dynamic genes (**Extended Data Fig. 3c**). Comparing wave-specific gene signatures between these two species revealed that *Acomys* astrocytes at birth express mature-like programs characteristic of *Mus* P7-P14 astrocytes (**Extended Data Fig. 3d, e**).

Given that *Acomys* cortex displays mature-like features at birth while *Mus* undergoes protracted postnatal maturation, chronological age matching (*e*.*g*., P0 to P0) does not align equivalent developmental stages between species. To enable a stage-matched molecular comparisons, we constructed an integrated pseudotime model using ordinal regression^22^ (**Fig. 3e, Methods**). Neuronal cells (both glutamatergic and GABAergic) from both species were ordered along a shared developmental trajectory. In both *Mus* and *Acomys*, glutamatergic and GABAergic populations progressed along a continuous pseudo temporal axis consistent with their developmental stage, with *Acomys* cells at birth occupying more advanced pseudotime positions than chronologically matched *Mus* cells (**Fig. 3e**). Earth Mover’s Distance (EMD) to the adult *Mus* reference—a measure of transcriptomic similarity—confirmed this asynchrony (**Fig. 3f**). In *Mus*, both glutamatergic and GABAergic neuronal populations showed gradual reduction in EMD from embryonic stages through postnatal development, reaching adult-like states by P15. In contrast, *Acomys* populations exhibited steeper EMD decline during pre/perinatal stages, with P0-P7 timepoints already approaching the adult *Mus* stage. Thus, the molecular maturation trajectory unfolding postnatally in Mus proceeds prenatally in Acomys.

To determine whether transcriptional trajectories that unfold postnatally in *Mus* are altered or simply shifted prenatally in *Acomys*, we examined developmental gene expression dynamics along pseudotime (**Fig. 3e-j, Methods**). Gene expression waves were computed separately for each species and cell type, allowing identification of dynamically expressed genes (**Extended Data Fig. 4a, b**). This approach revealed that an important fraction of gene expression is shared between the two species (**Fig. 3g**), with high correlation values supporting conserved dynamic expression. This conservation is largely shared across neuronal subtypes (**Fig. 3h**), suggesting that the dynamic transcriptional programs progress similarly in the two species across cell types. To classify the different waves of dynamic gene expression during maturation, we computed a UMAP-based clustering and identified two main groups: activated and repressed genes (**Extended Data Fig. 4d, Methods**). Highly correlated genes were predominantly activated over developmental time, with a smaller subset showing progressive repression (**Fig. 3i**). Activated gene programs were significantly enriched for biological processes associated with synaptic signaling and nervous system development, while repressed gene programs were associated with brain and central nervous system development (**Extended Data Fig. 4c, Methods**).

Because birth in *Mus* coincides with the onset of neuronal maturation, postnatal experience has been viewed as a driver of cortical maturation programs. To test whether birth timing itself is associated with major transcriptional transitions in *Mus*, we identified differentially expressed genes (DEGs) between sequential developmental stages (**Methods**). Surprisingly, for both glutamatergic and GABAergic neurons, a lower number of DEGs was found between prenatal (E17) and postnatal stages (P1) compared to later stage comparisons (**Fig. 3j** top, **Extended Data Fig. 4e**). This suggests that birth itself does not contribute to maturation-related changes in gene expression. Furthermore, perinatal gene expression changes in *Mus* (E17 vs P1 DEGs) show similar expression dynamics during prenatal stages in *Acomys* (**Fig. 3j** bottom, **Extended Data Fig. 4f**), demonstrating that birth does not constitute a major transcriptional transition in cortical differentiation programs. Together, these analyses show that, transcriptionally, cortical maturation programs are largely conserved between *Mus* and *Acomys*, but unfold irrespective of birth timing.

### Behavior-relevant somatosensory circuit assembly occurs before birth in *Acomys*

The presence of fully elongated whiskers, together with substantial morphological, electrophysiological and molecular maturation suggest that functional *Acomys* somatosensory circuits may mature to a significant extent before birth. A classical readout of rodents’ somatosensory circuit maturation is the formation of the primary SSp barrel field. Barrels correspond to the organizational unit of thalamocortical axon (TCA) terminals and L4 spiny stellate neurons (SSN), with each barrel receiving input from a single whisker from the contralateral whisker pad. In *Mus* newborns, TCAs form a diffuse band in SSp L4, which then cluster into discrete barrels between P2 and P4, and finally consolidated until P15 through activity-dependent refinement^4^. Strikingly, *Acomys* instead display a fully formed barrel field at birth (**Fig. 4a**). To determine when the barrels emerge during *Acomys* development, we examined TCA organization in *Acomys* at different embryonic timepoints. At E27.5, VGLUT2-positive TCAs form a diffuse layer in presumptive L4. However, by E29.5—corresponding to *Mus* P4—TCAs are already organized into discrete barrels (**Fig. 4b**), indicating that anatomical barrel formation occurs largely before birth.

**Figure 4.**
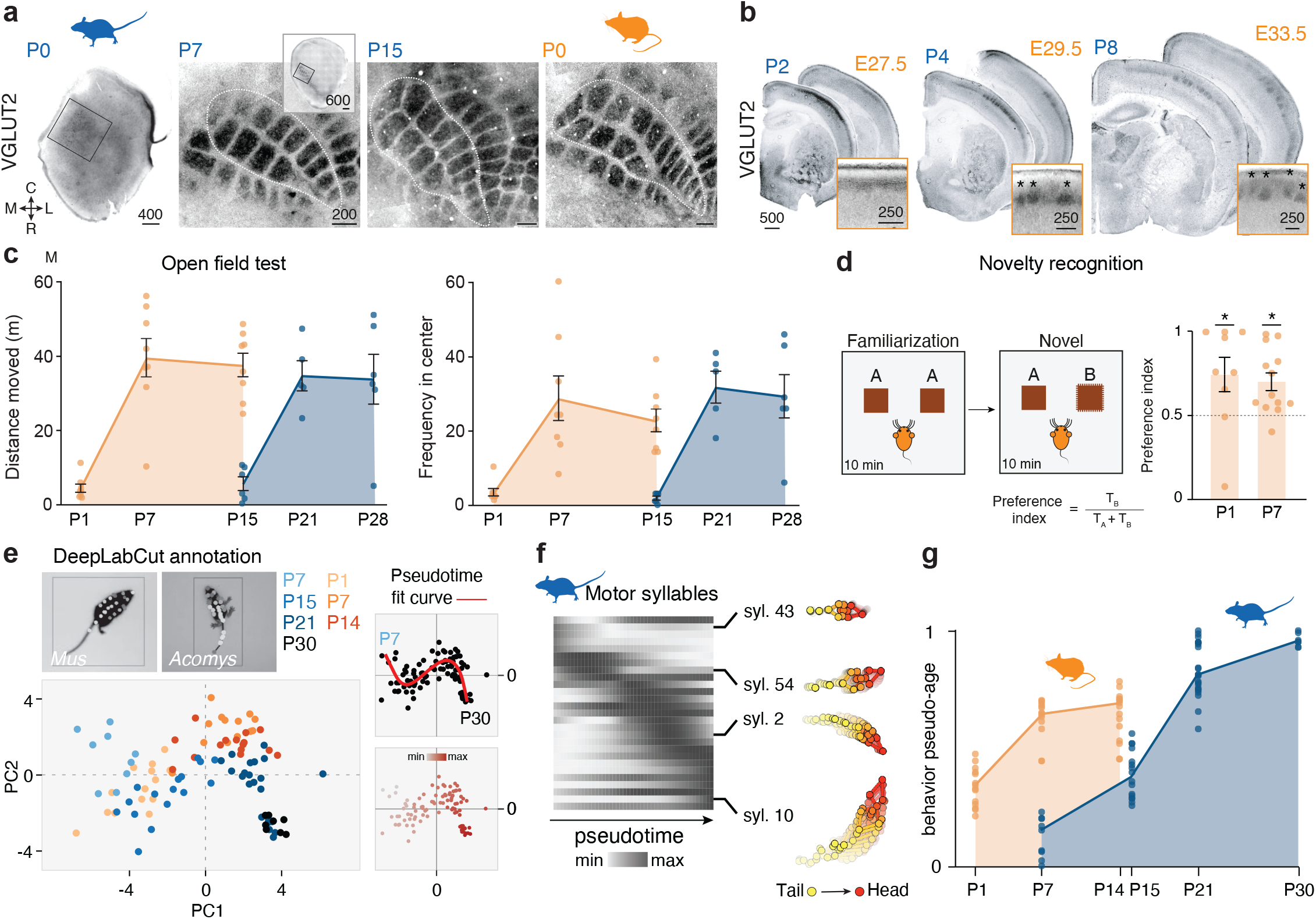
Circuit and behavioral maturity in *Acomys*. **a**, VGLUT2 immunofluorescence images of P0, P7 and P15 *Mus* and P0 *Acomys* flattened cortices. **b**, VGLUT2 immunofluorescence images of P2, P4 and P8 Mus and E27.5, E29.5 and E33.5 *Acomys* coronal brain sections. **c**, Distance moved (in meters, left) and of frequency to the center of the arena (right) during Open field test at P1, P7 and P15 in *Acomys* and P15, P21 and P28 in *Mus*. Lines indicate mean trajectory. **d**, Schematics of novelty recognition test (left) and bar plot of preference index at P1 (one sample t-test against chance level 0.5, p-value: 0.0442) and P7 (one sample t-test against chance level 0.5, p-value, 0.0025) in *Acomys* (right). **e**, Principal Component Analysis (PCA) plot of motor behavior in *Acomys* at P1, P7 and P14 and *Mus* at P7, P15, P21 and P30. Dots are color-coded per species (color) and ages (shades). Right: PCA plot of curve fitting (top) and pseudotime values (bottom) along the curve (color-coded). **f**, Heatmap representation of motor syllables across pseudotime in *Mus*, with example syllables highlighted on the right, where color code indicates the tail to head side of the animal. h, Behavioral pseudo-age values for *Acomys* and *Mus* at respective timepoints. Lines indicate mean trajectory. Abbreviations: P, postnatal day; E, embryonic day; PC, principal component. Points represent individual animals in c-g. Scale bar units: µm.

Concomitant with barrel formation, SSN in *Mus* undergo two morphological changes: their basal dendrites polarize toward the barrel center, and their apical dendrites retract^43^. To examine whether these morphological transformations also occur before birth in *Acomys*, we performed whole-cell recordings and morphological reconstructions of SSN in *Mus* (P7-P15) and *Acomys* (P1-P15). In *Mus*, SSN displayed polarized basal dendrites and lacked apical dendrites from P7 onwards (**Extended Data Fig. 5a**). In P1 *Acomys*, SSN showed strong basal dendrite polarization toward barrel centers but retained prominent apical dendrites (**Extended Data Fig. 5b**). Apical dendrite retraction began around P7 and was complete by P15 in *Acomys*. This reveals a temporal dissociation between dendrite polarization and apical dendrite retraction in this species. Despite this morphological difference, electrophysiological properties of *Acomys* P1 SSN closely resembled *Mus* P7 SSN, and both species showed similar maturation trajectories in PCA analysis (**Extended Data Fig. 5c**).

Active exploration of the environment requires sensory processing together with coordinated movement. We therefore assessed open-field exploration and novel object discrimination, behaviors that emerge progressively during postnatal *Mus* development^44^. Strikingly, unlike newborn *Mus*, newborn *Acomys* displayed increased locomotor activity and center exploration in the open-field arena (**Fig. 4c**), and discriminated novel from familiar objects (**Fig. 4d**). Thus, sensorimotor behaviors that emerge postnatally in *Mus* are already present at birth in *Acomys*.

To quantify this precocious sensorimotor maturation more comprehensively, we performed high-resolution behavioral tracking of *Mus* and *Acomys* pups across postnatal development using high-quality video recordings of both species (**Methods**). We trained a Motion Sequencing (MoSeq) model^45^ to extract movement “syllables”, *i*.*e*. stereotyped behavioral motifs (**Supplementary information video 3**), from these recordings, capturing the full repertoire of spontaneous movements. PCA of motor syllable features revealed a shared developmental trajectory for both species (**Fig. 4e**), with conserved changes in movement complexity and coordination. To align these behavioral trajectories across species, we constructed a pseudotime model based on PCA features (**Fig. 4f, Methods**), revealing that *Acomys* P1 corresponds to *Mus* P15 in terms of sensorimotor maturity (**Fig. 4g**). Together, these findings show how sensorimotor functions that emerge early postnatally in *Mus* are already available at birth in *Acomys*.

## Discussion

Our results show that cortical maturation is not intrinsically tied to birth. By comparing *Acomys* to *Mus*, two related rodents with markedly different gestation lengths, but a conserved corticogenesis time-window, we find that major histological, morphological, electrophysiological, molecular and behavioral features of cortical maturation, unfold along similar trajectories, despite occurring before or after birth. Thus, precociality does not appear to rely on a fundamentally different cortical maturation program, or its timing, but rather reflects the prenatal displacement of conserved programs that, in *Mus*, unfold postnatally. Indeed, for cortical neurons maturation, birth does not appear to be a significant developmental landmark. For cortex, birth does not appear to be a significant developmental landmark, but rather an evolutionarily movable transition point placed at different positions along a conserved cortical maturation trajectory.

Despite its nearly two-fold longer gestation, Acomys undergoes corticogenesis over a time window similar to Mus. The additional prenatal time is therefore used not to produce more cortical neurons^46–50^, but to advance neuronal and glial maturation, thalamocortical circuit assembly and sensorimotor function before birth^10^. However, this does not imply that precocial prenatal maturation is activity independent. Rather, our findings potentially reflect prenatal activity-dependent processes, such as spontaneous activity or whisker contacts with the uterine wall. Indeed, active whisking and novel object discrimination at birth suggest that some degree of prenatal sensorimotor processing is already present. Notably, even in *Mus*, whisker-related hindbrain “barrelettes” are already present at birth^51^, indicating that aspects of somatosensory circuit patterning emerge before postnatal experience^52^.

Although the behavioral paradigms that we have used were developed in *Mus* and may not be identically interpretable in *Acomys*, the consistency of these results with those obtained at both molecular and cellular levels suggests that the overall precocious maturation enables precocious behavioral function. Immaturity at birth may confer the advantage of allowing cortical circuits to adapt to specific sensory and environmental conditions during postnatal development. By contrast, the ecological constraints faced by *Acomys*, including the need for early mobility and sensory competence in harsh desert environments, may have favored greater developmental maturity at birth, potentially at the cost of a reduced capacity for experience-dependent circuit refinement. Therefore, it remains to be tested to what extent postnatal experience-dependent refinement occurs in *Acomys*, and how this relates to *Mus* in the context of their species-specific behavioral repertoires.

Across evolution, birth can occur at different stages of conserved maturation programs. Within a species, however, birth and cortical maturation are normally aligned. Prematurity disrupts this alignment, which is thought to contribute to altered cortical connectivity and adverse neurodevelopmental outcomes^53,54^.

Finally, the limited effect of birth itself on cortical maturation may not be entirely surprising, as the cortex remains relatively remote from immediate peripheral and homeostatic transitions occurring at birth. Developmental divergence associated with intra-vs. extrauterine maturation may be more pronounced in brain regions more directly involved in physiological adaptation and survival, such as the hypothalamus (hormonal regulation) or brainstem (e.g. breathing). Extending this comparative framework across additional brain regions and species will help to delineate how conserved developmental programs are aligned relative to birth, and whether longer prenatal maturation limits later postnatal circuit plasticity and experience-dependent refinement.

## Acknowledgments

We thank D. Valloton, K. Ascencao, C. Rummel, and A. Liechti for RNA-seq data generation. The RNA-seq libraries were sequenced at the Lausanne Genomic Technologies Facility. We thank the iGE3 genomics facility of the University of Geneva, the HCNP and the Wyss Imaging platforms of the Campus biotech in Geneva for technical support. We thank all the members of the Jabaudon and the Tzika laboratory for their comments on the manuscript and for constructive exchanges during the project. ACT is supported by the Swiss National Science Foundation (310030_204466), the HFSP (RGP0037/2022), the Ernst and Lucie Schmidheiny Foundation (10_2023), the Fonds Général de l’Université de Genève (23_28), and the Emile Plantamour Fund (2024/12). The Jabaudon laboratory is supported by the Swiss National Science Foundation, the Société Académique de Genève FOREMANE Fund, and the European Research Council; I.M was supported by iGE3 PhD Salary Award 2023 (no. FNS ME12454); S.F. was supported by the Swiss National Science Foundation (Ambizione grant PZ00P3_193349); R.B. and J.Z. were supported by the Swiss National Science Foundation (Ambizione grant: PZ00P3_201995); CB was supported by the Swiss National Science Foundation (310030-212219); SRP was supported by EMBO (ALTF 349-2020).

## Authors contributions statement

A.P. and C.B. designed behavioral experiments; A.P. performed behavioral experiments on *Mus* and analysis of both *Mus* and *Acomys* behavioral experiments; S.V. collected data of birth-dating experiments which was analyzed by S.R.P and I.M.; F.M., A.D. and A.C.T. handled the *Acomys* colony and performed behavioral assessments for all *Acomys* experiments; A.C.T provided the samples for generation and annotation of the *Acomys* reference genome. J.Z. and R.B. performed the bioinformatic analysis on astrocytes maturation; J.P. assisted with bioinformatic analyses; S.F. performed electrophysiological experiments; S.V. and E.K. performed initial histology experiments. I.M. and S.R.P designed and conducted experiments, analyzed and interpreted the data; I.M., D.J. and S.R.P. conceived the project and wrote the manuscript. A.P., S.F., S.V., J.P., F.M., A.D., J.Z., R.B., E.K., C.B. and A.C.T. corrected and gave feedback on the manuscript.

## Competing interests’ statement

The authors declare no competing interests.

## Methods

### Animal strains and husbandry

Wild type mice were obtained from Charles River Laboratory. *Acomys* were housed and bred at the LANE animal facility running under the veterinary cantonal permit no. 1008. All the experimental procedures described here were conducted in accordance with the Swiss laws and previously approved by the Geneva Cantonal Veterinary Authority (GE24/33145 and GE205 approved protocol). Swiss mouse husbandry was done at the institutional animal facilities under standard 12h:12h light:dark cycles with food and water ad libitum. Embryonic day (E) 0.5 (overnight mated females) was established at time of detection of vaginal plug. In this study we indistinctively analyzed males and females. Spiny mice are kept under a 14h:10h light:dark cycle with food and water available ad libitum. A vaginal plug is not formed in spiny mice. Females were bred for two consecutive days, and the pregnancy was verified by palpation at the appropriate stage.

### Immunohistochemistry

The same procedure was used to collect *Mus* and *Acomys* samples. Animals were perfused with 4% PFA/PBS and brains first fixed overnight in 4% PFA/PBS at 4 °C, then cryoprotected at 4 °C in 20% sucrose (Sigma, cat# S0389) in PBS until sinking. Brains were then embedded in Tissue-Tek O.C.T. (Sakura Finetek, no. 4583) and preserved at −80 °C until cryosectioning at 60 µm (Leica CM3050). For free-floating immunostaining, sections were first washed in PBS with 0.1% Triton X and then incubated for 2 h at room temperature in blocking-permeabilizing solution containing 3% bovine serum albumin and 0.3% Triton X-100 in PBS, with subsequent incubation for 2 days at 4 °C with primary antibodies: CTIP2 (abcam #ab18465), Cux1 (Proteintech #11733-1-AP), MBP (abcam #ab7349), L1CAM (Merck Millipore # MAB5272), RORB (Bio-Techne #PP-N7927-00), VGLUT2 (Merck Millipore #AB2251), S100β (abcam #ab41548), BrdU (Life Technologies #B35128). Sections were then rinsed three times in PBS with 0.1% Triton X and incubated with the corresponding Alexa-conjugated secondary antibodies (1:500, Invitrogen) for 2 h at room temperature. The following Invitrogen secondary antibodies were used (Thermo Fisher Scientific, anti-mouse 488 #A21292, anti-mouse 546 #A10036, anti-mouse 647 #A31571, anti-rabbit 488 #A21206, anti-rabbit 647 #A31573, anti-guinea pig 488 #A11073, anti-guinea pig 647 #A21450, anti-rat 488 #A21208, anti-rat 647 #A48272). Nuclei were stained with Hoechst (Invitrogen, no. H3570)/PBS during secondary antibody incubation. Images were acquired with Tissue Scope and Nikon A1X with 20X objectives.

### Flatten cortices

The same procedure was used to collect *Mus* and *Acomys* samples. Animals were perfused with 4% PFA/PBS and brains fixed for 2 hours at room temperature. Following fixation, individual hemispheres were flattened using glass slides (Menzel Gläser) and fixed for 3hrs at room temperature. For sectioning, the flatten cortices were included in 4% agarose (Sigma, cat #A9539) prior to slicing at 70µm using Leica VT1000 S vibratome. For immunostaining, the same procedure as described in the dedicated section was used. The following primary antibody was used: VGLUT2 (Merck Millipore #AB2251).

### Neurogenesis birth-dating injections and immunofluorescence

At the timepoint of interest, *Acomys* females were injected intraperitoneally with 10mg/ml BrdU (Thermo Fisher #H27260) or with 7.5mg/ml EdU (Thermo Fisher, #A10044), dissolved in 0.9% NaCl, for a dosage of 50mg/kg of animal weight. To minimize the use of animals, the following injections were done in the same animals: E16.5 EdU + E24.5 BrdU; E19.5 BrdU + E22.5 EdU; E20.5 BrdU + E23.5 EdU. The tissue was collected for immunohistochemistry at P 21 for all animals. For BrdU staining, free floating sections were washed 3×10 min in PBS at room temperature, then denatured prior to blocking by incubating them in 2N HCl at 37°C for 30 min and washed in PBS 2×15min. The sections were then incubated 1h at room temperature in blocking solution (10% normal horse serum, 0.1% Triton-X 100 in PBS) and then incubated at 4°C overnight with primary antibody diluted in the same blocking solution. Sections were then washed in PBS 3×15 min and incubated 2h with the secondary antibody diluted in blocking solution. After washing again 3×15 min with PBS, sections were mounted with Sigma Fluoromount. For EdU staining, the Click-IT chemistry was done following the manufacturer’s instructions (Invitrogen). Sections were then incubated with DAPI (1:1000 in PBS) for 15 minutes at RT, washed in PBS for 10 min and mounted with Sigma Fluoromount.

### Electrophysiological recordings

Three-hundred-micrometer-thick acute coronal slices were prepared from *Mus* and *Acomys* brains at different developmental stages. Slices were kept for at least 30 min in modified artificial cerebrospinal fluid (aCSF) at 33°C (87mM NaCl, 2.5 mM KCl, 7mM MgCl2, 0.5mM CaCl2, 1.25mM NaH_2_PO_4_, 25mM NaHCO3 and 5 mM glucose), supplemented with 1mM kynurenic acid (Sigma K3375) and oxygenated with 95% O2 and 5% CO2 which was gradually replaced by aCSF used for patch-clamp recording (125 mM NaCl, 2.5 mM KCl, 1mM MgCl2, 2.5mM CaCl2, 1.25 mM NaH_2_PO_4_, 26 mM NaHCO3 and 11 mM glucose, oxygenated with 95% O2 and 5% CO2). The slices were transferred to the recording chamber, submerged, and continuously perfused with aCSF. Patch-clamp recordings were performed on cells from different apico-basal levels of SSp, targeting L2/3, L4 and L5b neurons. The internal solution used for the experiments contained 140mM potassium methansulfonate, 2 mM MgCl2, 4 mM NaCl, 0.2 mM EGTA, 10 mM HEPES, 3mM Na2ATP, 0.33mM GTP and 5mM creatine phosphate, 0.3% biocytin; pH 7.2; 295 mOsm. In voltage-clamp configuration, the voltage was clamped at -60mV. Access resistance was monitored by a hyperpolarizing step of 14 mV. Ih current was calculated following a hyperpolarization step of -40mV for 500ms. The neuron was then placed in current clamp mode, and the resting membrane potential was monitored every 10 s and averaged for 5 consecutive acquisitions. Four steps of 50, 100, 200 and 400 pA were applied for 500 ms. During the recording, neurons were passively filled with biocytin and at the end of the recording, the patch pipette was slowly retracted from the cell membrane to obtain an outside-out patch to maintain the integrity of the plasma membrane for further neuron morphology analysis. Fifteen minutes after the end of the recording, slices were fixed with 4% paraformaldehyde at 4°C overnight and incubated with Alexa 647 coupled-streptavidin (Invitrogen S21374, 1:500 in PBS-10% tween) for 6 h at room temperature. Sections were then washed with PBS before mounting. Analyzed neurons were recorded from different animals per species (**Supplementary Information Table 2**).

### Morphological reconstructions and analysis

Biocytin filled imaged neurons were imported in Imaris and neuronal morphology was manually reconstructed using the filament tool.

### Acomys dimidiatus genome sequencing and annotation

High molecular weight (HMW) DNA was extracted from kidney samples using the Quick-DNA HMW MagBead Kit (Zymo Research, D6060). Genome sequencing was outsourced to Novogene^®^. Illumina (173.10X) and Nanopore (75.46X) sequencers were used to obtain high quality short and long reads (total coverage: 248.56X). Wtdbg2 (https://github.com/ruanjue/wtdbg2) was used for the assembly, and the contigs were polished with Racon for Nanopore reads and Pilon for the Illumina reads. Genome completeness was evaluated by BUSCO (https://busco.ezlab.org), which showed that 91.9% of 13798 single-copy orthologs are found in the genome as complete genes, and similarly with CEGMA (http://korflab.ucdavis.edu/datasets/cegma/), which showed 93.95% of completeness.

Read mapping rate (98.91%) and coverage (99.39%) were also confirmed by Burrows-Wheeler aligner assessment (https://bio-bwa.sourceforge.net). For genome annotation, RNA sequencing samples were generated. We collected samples from brain, testis, ovary, cerebellum, kidney, liver and heart from either male or female samples. RNA was extracted using the RNeasy protocol from Qiagen. RNeasy Micro columns (no. 74004: Qiagen) and RNeasy Mini columns (no. 74104; Qiagen) were used to extract RNA from small (<5 mg) and larger (>5 mg) samples, respectively. The tissues were homogenized in RLT buffer supplemented with 40 mM dithiothreitol (DTT) or QIAzol. RNA quality was assessed using the Fragment Analyzer (Advanced Analytical); RQN values were greater than 6.7 for all samples. The RNA-seq libraries were prepared using the TruSeq Stranded mRNA LT Sample Prep Kit (Illumina) and sequenced on the HiSeq 2500 platform. We obtained 24.1 to 50 million 100-bp strand-specific single-end reads per library. We also extracted total RNA from the dorsal skin of an embryo using the Direct-zol RNA MiniPrep (Zymo Research, R2050) and obtained a RIN of 9.7. 100 bp paired-end TruSeq Stranded mRNA libraries were prepared and produced 21 million reads. Annotated genome and RNA sequencing samples will be released from embargo upon manuscript acceptance.

### Single-nuclei RNA sequencing of S1 Acomys dimidiatus samples

Somatosensory cortices (S1) were collected for E25 embryos (n=2), P0 (n=3), P7 (n=3), P14 (n=3) and P113 (n=1) *Acomys*. Brains were first micro-dissected, and 300-400 µm-thick coronal slices were prepared using Leica VT1200S Vibratome. After, the region of interest was micro-dissected under stereomicroscope (Nikon) ice-cold HBSS 1x (Gibco, cat#14025092). The micro-dissected tissue was preserved at -80°C until use for the single nuclei suspension, which was obtained through douncer homogenization (Sigma D8938) using an adaptation of the protocol for the Nuclei EZ buffer kit (Sigma, NUC101). For dissociations, after being thawed on ice, the tissue was resuspended in 2mL of ice-cold extraction buffer (Nuclei EZ prep buffer) and dounce-homogenized on ice until disintegration (5 to 15 pestle A and B). After incubation for 5 minutes in 4ml EZ buffer total, the extracted nuclei were collected by centrifugation (500 x g at 4°C, 5 minutes). The extraction procedure was repeated once, before performing a washing step, in which nuclei were resuspended in washing buffer composed of 1% BSA in PBS, with 50U/ml of SuperASE-In (Thermo Fisher, cat# AM2696) and 50U/ml of RNasin (Promega, cat# N2611), after centrifugation. Finally, the nuclei were resuspended in 1mL of washing buffer and filtered through a 30μm strainer into a new low-binding 1.5mL tube. 20-40k nuclei were loaded into the 10x genomics Chromium Next GEM Single Cell 3’ Kit v3 (P7, P14) or GEM-X Universal 3’ Gene Expression (E25, P0, Adult) according to the manufacturer’s instructions.

### Mapping of snRNA-seq data

*Acomys dimidiatus* genome was annotated using EGAPx (https://github.com/ncbi/egapx) using the novel genome assembly and RNA-seq samples (described in the previous section). The reference transcriptome was generated using Cellranger 8.0.1 mkref.

All single-nucleus transcriptomics data generated using 10x Genomics 3’ GEX kits were mapped using Cellranger 8.0.1 *count*.

### Analysis of snRNA-seq S1 dataset

#### Quality controls, annotation, integration

Low quality cells (<800 genes and >2.5% mitochondrial RNA) and doublets were removed during the initial step of quality control, done for each dataset separately. All count matrices were normalized and scaled using Seurat dedicated functions. PCA was then calculated on the top 2000 most variable genes and UMAP projections were built on PCA and k-nearest neighbors for each dataset separately. Annotation was done using the online tool “MapMyCells” (https://knowledge.brain-map.org/mapmycells/process/).

After annotation, S1 Acomys datasets were merged and harmonized using Seurat, and contaminations of non-cortical cells were removed. Subsequently, the Acomys S1 dataset was saved in .h5ad format for integration using SysVI methodology (scvi-tools package in python). For the Mus reference dataset, E17, P1, P7 and P14 datasets were taken from^39^ and Adult S1 cells were taken from Langlieb, J. et al. (NeMO ID: nemo:dat-y5zxh0y)^40^. The integration was done with SysVI cVAE model, trained with KL weight of 1, using 89 epochs, the species as batch-key and the individual samples as categorial covariate keys. The plots in Figure 3 and Extended Data Figures 2-4, were generated using Seurat dedicated functions and ggplot2.

#### Pseudotime analysis

Pseudotime was built using ordinal regression models on Mus and Acomys data from *bmrm* package as done previously^22^. Pseudotime values were centered by subtracting the median pseudotime of mouse P1 cells. Analyses were restricted to glutamatergic and GABAergic neuronal populations. To quantify differences in developmental state relative to adulthood, we computed the Earth Mover’s Distance (EMD, Wasserstein distance) between pseudotime distributions. Single-cell metadata were extracted and grouped by species and subclass identity. For each subclass in each species, we calculated the one-dimensional Wasserstein distance between its pseudotime distribution and the corresponding *Mus* Adult reference distribution using the *wasserstein1d* function from the *transport* R package. Results were represented using ggplot2 in R. For astrocytes pseudotime analysis, the same methodology was applied on *Mus* publicly available developmental datasets^55–58^.

#### Developmental gene wave analysis and cross-species comparisons

To model gene expression dynamics along pseudotime, we applied Gaussian kernel smoothing to log-normalized expression values. For each species and neuronal subclass independently, pseudotime was truncated to a fixed range and discretized into 25 evenly spaced bins. For each gene, smoothed expression across bins was computed using a Gaussian kernel (σ = 0.3), generating a matrix of genes × 25 pseudotime points representing the gene’s developmental wave. Genes with insufficient dynamic range (max - min ≤ 0.1 across bins) were excluded from downstream wave embedding analyses. Subclasses with discontinuous pseudotime coverage were omitted. For visualization, wave profiles were min–max normalized per gene.

#### Gene waves analysis

To compare wave profiles across genes, the 25-bin expression vectors were embedded using UMAP with correlation distance. Genes were clustered in UMAP space using Louvain community detection on a k-nearest neighbor graph. Clusters were annotated based on the average standardized wave profile. Cluster-level average wave profiles were computed after z-scoring each gene across bins. To identify biological processes associated with conserved developmental dynamics, Gene Ontology (GO) enrichment analysis was performed on genes exhibiting highly conserved wave profiles between species (pearson correlation > 0.75). GO Biological Process annotations were obtained from curated *Mus musculus* pathway databases using *clusterProfiler* package.

#### Behavioral assessments

To conduct the behavioral characterization, *Mus* and *Acomys*, male and female of 1 day to 30 days of age were used. All behavioral assays were performed during the light phase of the light/dark cycle, in a soundproof cabinet or room with fixed illumination (25 Lux, unless specified differently) and with controlled humidity (40–50%) and temperature (22–24 °C), which were maintained consistently throughout all experiments to ensure validity. All animals were habituated to the experimental room for 1h prior to the beginning of the experiments. Every test session was video-tracked (*Mus*: Basler acA 1300-60gm camera with Kowa manual c-mount 4.4-11mm/F1.6 1/1.8” lens; *Acomys*: Sony FDR-AX53 16.6MP 4K Ultra HD Handycam Camcorder) and recorded at 25fps (using Ethovision XT from Noldus for *Mus*), which provided an automated recording of the position of the nose, body center, and tail points. This information was used to derive analysis of locomotion, exploratory behavior, and position in the arena.

#### Novel stimuli recognition test

The novel texture recognition test was conducted in a 30 × 30 × 30 cm arena under 25 lux illumination. The protocol consisted of three consecutive phases, each lasting 10 minutes. First, animals were habituated to the empty arena. This was immediately followed by the familiarization phase. After completion of this phase, animals were returned to their home cage for a 20-minute interval, after which the novel texture discrimination phase was performed. Two pairs of objects were used in the test, consisting of custom made, 3D printed cubes (3 cm). Both pairs were covered with orange sandpaper (Wolfcraft), with one pair using fine granularity paper (P120) and the other using coarse sandpaper (P40). The object pairs were alternated between animals to control for potential bias. During the familiarization phase, the two objects were placed at opposite sides of the arena. The animal was placed at the center of the arena and allowed to explore freely for 10 minutes. At the end of this phase, the mouse was returned to its home cage. After a 20-minute delay, the novel texture recognition phase began. One of the two objects was replaced with a novel object (the object to be replaced, left or right, was randomized between animals), while the other object remained the same (familiar object). Individuals were again placed in the center of the arena and allowed to explore freely. Recordings of all trials were processed using EthoVision XT, which automatically tracked key parameters such as distance moved (cm) and velocity (cm/s) for motor evaluation. Manual scoring was performed to record the time spent interacting with each object during both test phases. The arena was cleaned with 70% ethanol between trials. Key parameters of investigation, including preference indices and total time interacting with provided objects, were calculated for each phase from manual annotations. During the familiarization phase, the preference index was calculated as: preference index = [Total time spent interacting with object 1 /(Total time spent interacting with object 1 + total time spent interacting with object 2)]. The total time spent interacting with any object was calculated by summing the interaction times with both objects. Similarly, for the novel texture recognition phase, the preference index was calculated as: [Total time spent interacting with unfamiliar texture /(Total time spent interacting with unfamiliar texture + total time spent interacting with familiar texture)]. The total time spent interacting with any objects in the novel object recognition phase was also the sum of the times spent with both objects.

#### Open field test

The open field test was conducted to assess locomotor activity. The test was performed in a 50 × 50 × 50 cm arena (40 x 40 x 40 for *Acomys*) illuminated with approximately 160 lux. Each individual was placed at the center of the arena and allowed to explore freely for a 10-minute session, which was recorded for analysis. Key parameters, including distance moved (cm) and velocity (cm/s), were measured using Ethovision XT software. The arena was cleaned with 70% ethanol after each trial.

#### Motor behavior analysis

To achieve a refined characterization of motor behavior, animals were further analyzed using marker less pose estimation and unsupervised behavioral modeling. Videos acquired during the habituation phase of the novel texture discrimination test were used as input data (*Mus*: P7 = 12, P15 = 16, P21 = 23, P30 = 8; *Acomys*: P1 = 15, P7 = 13, P14 = 15). All videos were preprocessed to isolate the arena and remove background elements, thereby improving tracking accuracy. Pose estimation was performed using the DeepLabCut Model Zoo^59,60^, and keypoints corresponding to multiple body parts (including head, trunk, limbs, and tail segments) were extracted. The resulting keypoint data were used as input for unsupervised behavioral segmentation using Keypoint-MoSeq^45^. The model was trained on a representative subset of the dataset, including at least one example per strain and developmental time point. For downstream analysis, custom Python scripts were used. Only the most frequently occurring behavioral motifs were considered. PCA was performed using the following parameters: total distance travelled during the 10-minute session, frequency of the most represented motifs, and duration of these motifs.

#### Packages used and versions

The snRNA-seq datasets were analyzed using R statistical software (v4.4.1), SingleCellExperiment (v1.32.0), scater (v1.38.0), Seurat (v5.3.1.9999), ggplot2 (v4.0.0), transport (v0.15-4), tidyverse (v2.0.0), scDblFinder (v1.20.2), clusterProfiler (v4.18.2), patchwork (v1.3.2), bmrm (v4.4), dplyr (1.1.4), tidyr (v1.3.1), harmony (1.2.3), and using Python 3.11, scvi-tools SysVI (v1.4.0), Scanpy (v1.11.4), NumPy (v2.3.4), Pandas (v2.3.3), Matplotlib (v3.9.4).

#### Writing process using generative AI and AI-assisted technologies

The writing process used generative AI and AI-assisted technologies, with ChatGPT and Claude.ai used to improve specific sections of the paper. After using it, we reviewed and edited the content as needed and bear full responsibility for the content. Experimental procedures are presented in a narrative sequence for clarity; actual chronological order may differ from this presentation.

**Extended Data Figure 1.**
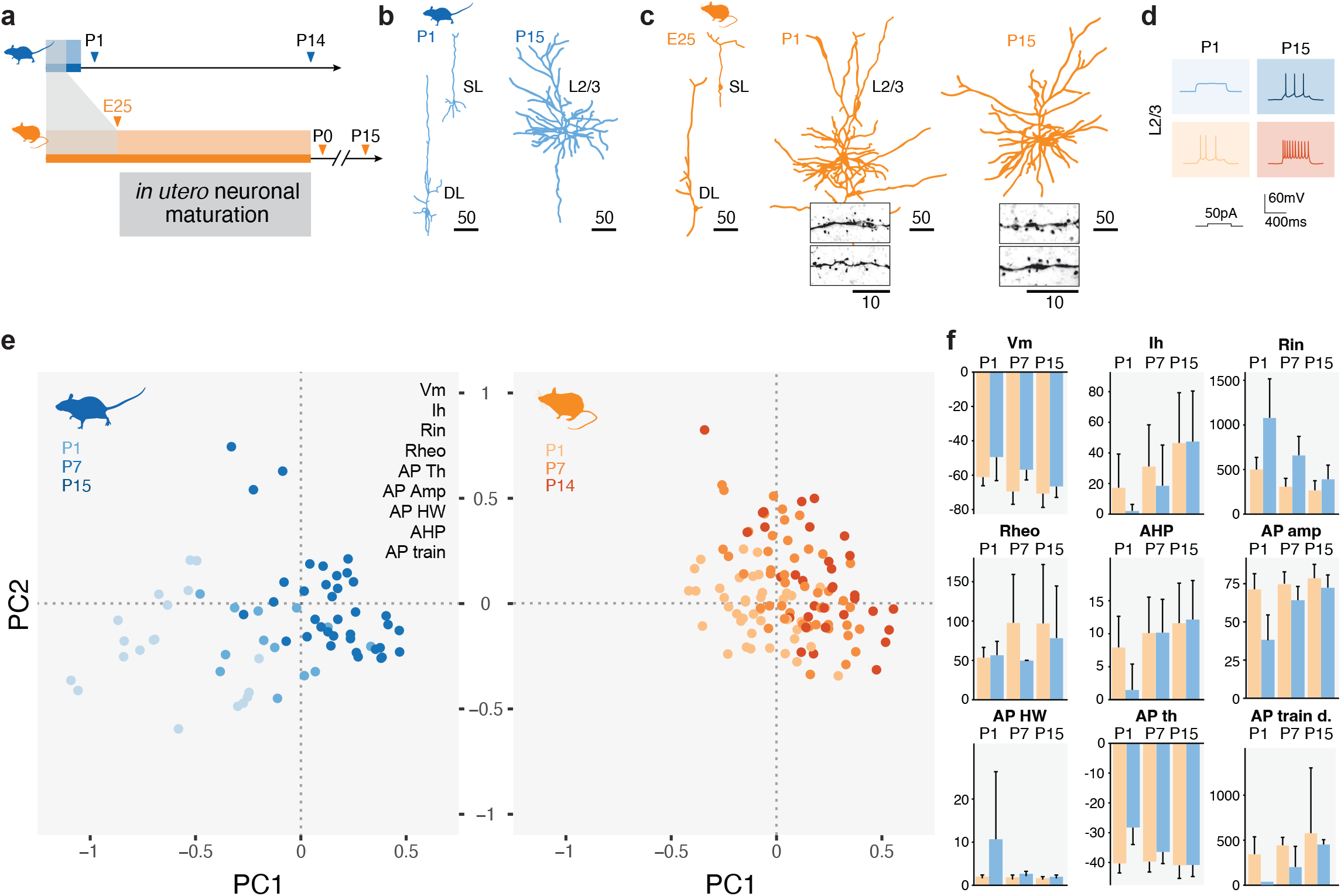
Characterization of morphological and electrophysiological features. **a**, Scheme of the developmental timelines. **b**, Example of reconstructed neurons in *Mus* at P1 (left) and P15 (right). **c**, Example of reconstructed neurons in *Acomys* at E25 (left), P1 (middle) and P15 (right). Insets include example of spines morphology in corresponding neurons. **d**, Example of traces of depolarizations in *Mus* (top) and *Acomys* (bottom), at P1 (left) and P15 (right). **e**, PCA plots of electrophysiological analysis in *Mus* (left) and *Acomys* (right) neurons across timepoints (color-coded). **f**, Bar plots of each individual electrophysiological parameter per species (color-coded) and across timepoints (columns). Scale bars are in µm. Abbreviations: PC, principal component; Vm, resting membrane potential; Ih, hyperpolarization-activated current; Rin, input resistance; Rheo, rheobase; AP Th, action potential threshold; AP amp, action potential amplitude; AP HW, action potential half-width; AHP, hyperpolarization potential; AP train, action potential train.

**Extended Data Figure 2.**
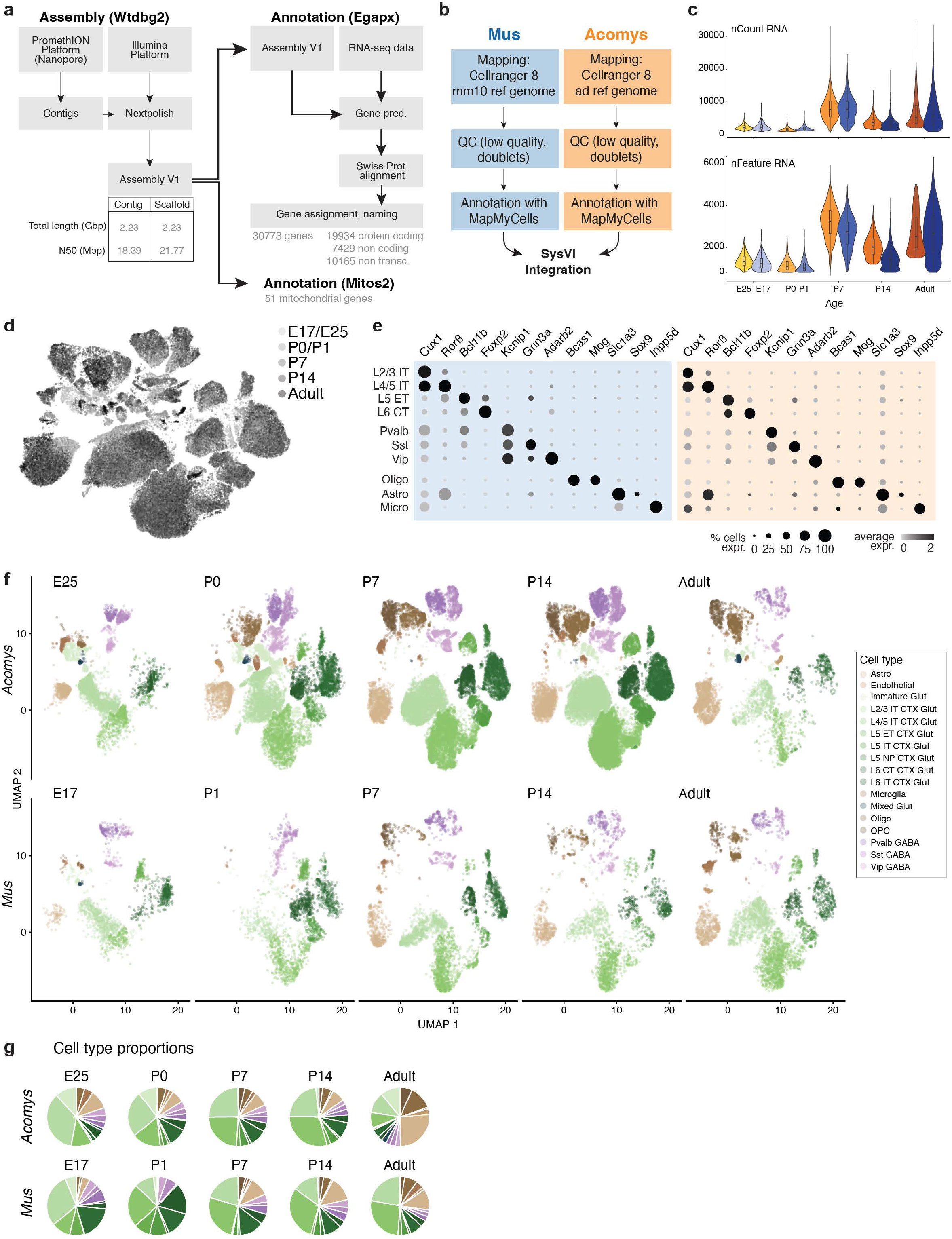
Mapping and integration of *Mus* and *Acomys* snRNA-seq data. **a**, Scheme of experimental and analytical strategy for the construction of the *Acomys dimidiatus* reference transcriptome. **b**, Scheme of analytical strategy to obtain mapped snRNA-seq data and integrated *Mus* and *Acomys* dataset. **c**, Violin plot of number of reads (nCount RNA) and number of genes (nFeature RNA) in *Mus* and *Acomys* (color-coded) across the different time points (columns). **d**, UMAP of integrated objects representing time points (color-coded). **e**, Dot plots for *Mus* and *Acomys* of canonical marker genes per cell type. The size of the dot represents the number of cells expressing the gene while the color indicates the expression level. **f**, UMAP of individual datasets for *Acomys* time points (top row) and *Mus* time points (bottom row), color-coded per cell type annotation. **g**, Pie charts per time points (columns) and species (rows) representing cell type relative proportions. Abbreviations: Astro, astrocytes; CTX, cortex; E, embryonic day; ET, extratelencephalically-projecting neurons; GABA, GABAergic neurons; Glut, Glutamatergic; IT, intratelencephalically-projecting neurons; Oligo, oligodendrocytes; OPC, oligodendrocyte precursor cells; P, postnatal day.

**Extended Data Figure 3.**
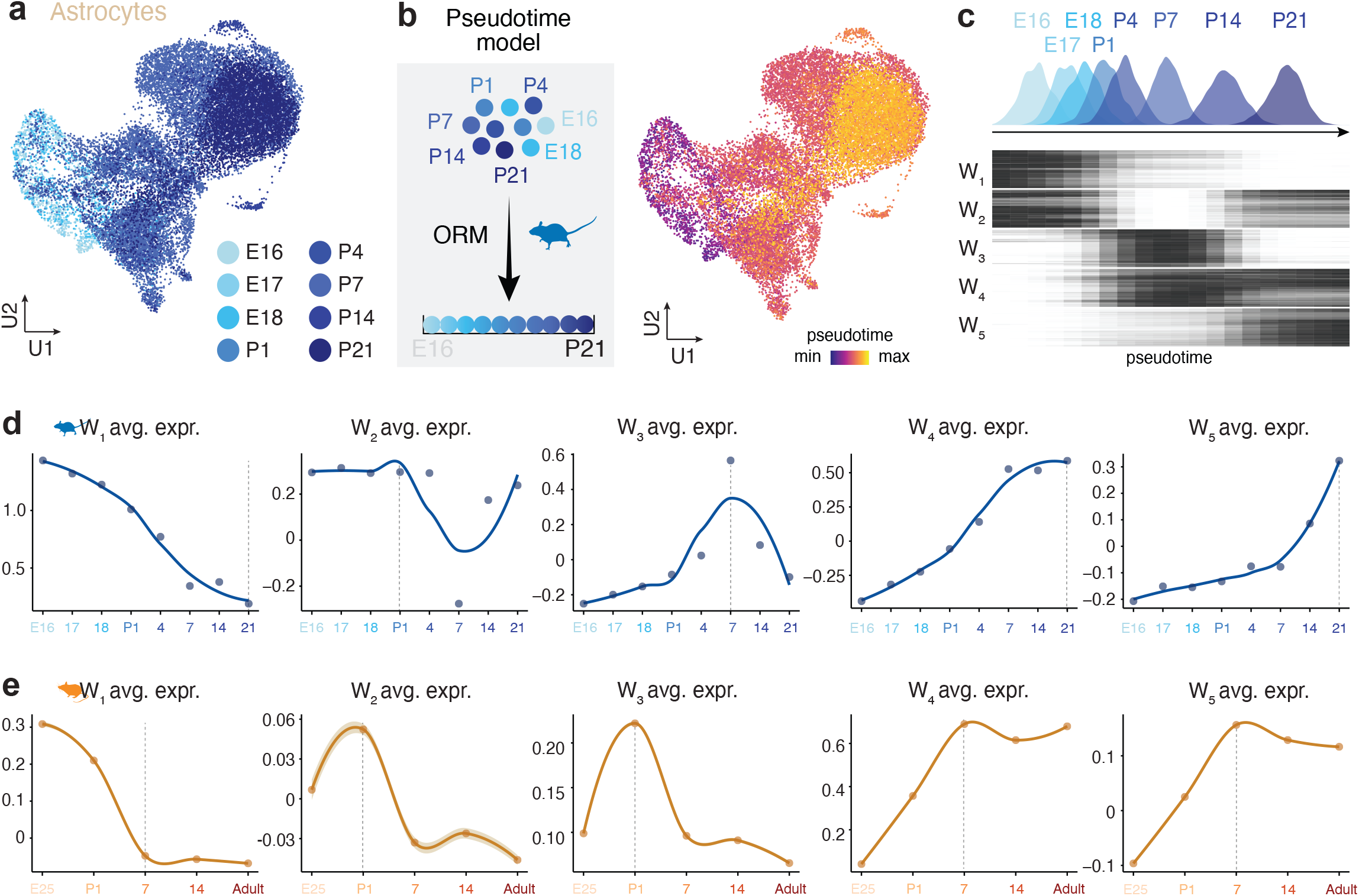
Temporal dynamics of astrocytes maturation across cortical development. **a**, UMAP visualization of integrated scRNA-seq datasets illustrating *Mus* cortical astrocytes development from embryonic day 16 (E16) to postnatal day 21 (P21). Cells are colored according to developmental stage. **b**, Schematic of the ordinal regression machine-learning (ORM) framework used to infer astrocytes pseudotime from developmental stage information (left). The model identified 500 genes predictive of astrocytes maturation, which were used to reconstruct a continuous trajectory from immature to mature astrocytes states (right). Cells are colored according to pseudotime progression. **c**, Heatmap showing the dynamic expression patterns of the 500 pseudotime-associated genes, organized into five temporal waves (W1–W5; 100 genes per wave) corresponding to sequential phases of astrocytes maturation. Density plots above the heatmap indicate the distribution of cells from each developmental stage along the inferred pseudotime axis. **d**, Average expression profiles of genes belonging to each temporal wave (W1–W5) across developmental stages in the *Mus* dataset shown in (a), illustrating the sequential activation and repression of distinct transcriptional programs during astrocytes maturation. **e**, Projection of the same wave-associated gene sets (W1–W5) onto the *Acomys* astrocytes developmental dataset generated in this study. Average expression profiles reveal broadly conserved maturation dynamics across species, with temporal waves peaking at earlier developmental stages in *Acomys* compared to *Mus* (dashed lines).

**Extended Data Figure 4.**
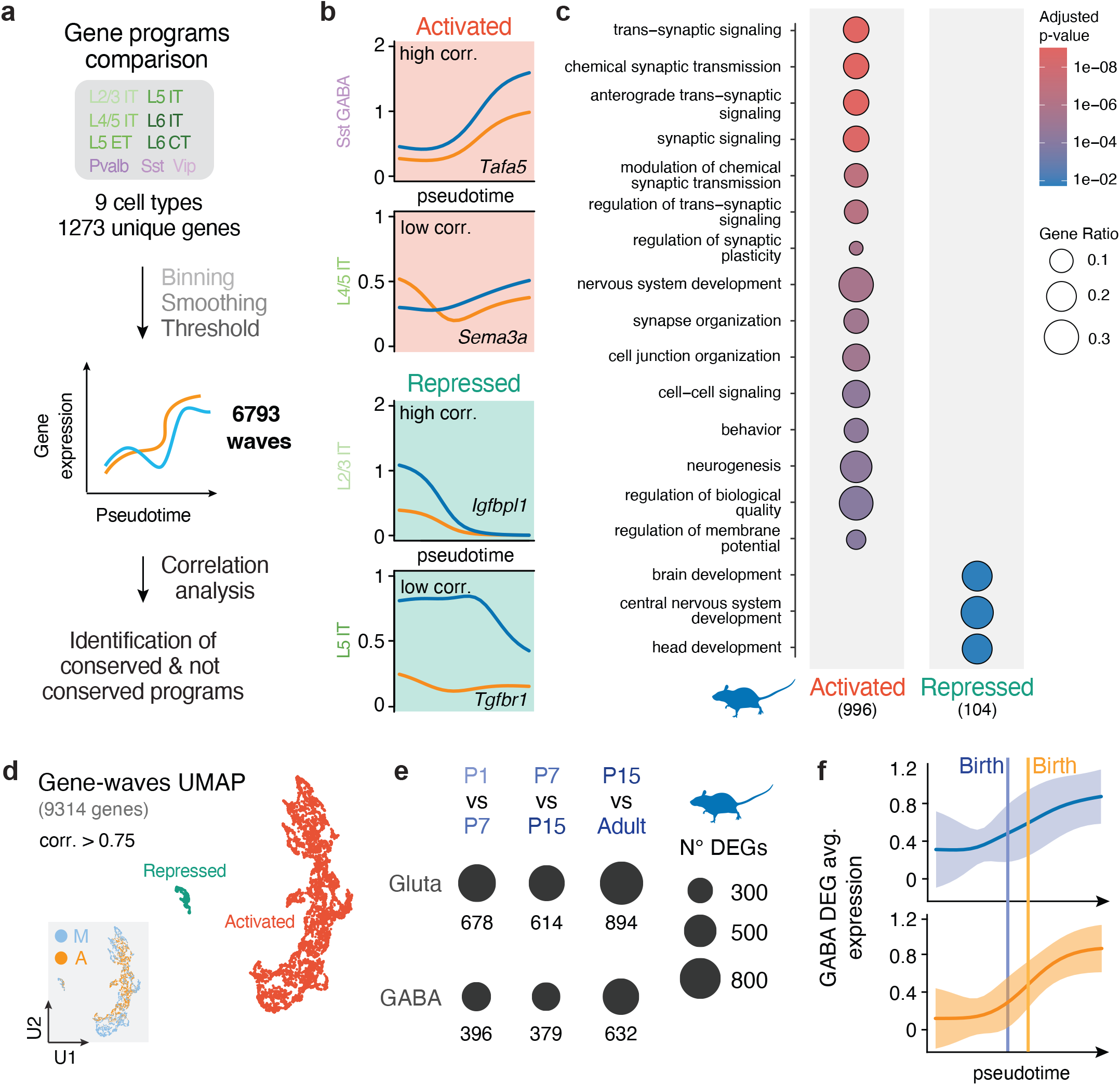
Gene wave analysis and birth-related gene expression. **a**, Schematics of gene waves analysis. **b**, Example of activated and repressed gene waves with either low or high correlation values. **c**, Dot plot of GO term biological process enrichment analysis results, for highly correlated (corr > 0.75) waves, for activated (left) and repressed (right) waves. The color of the dots represents the p-value, and the size of the dot represents the gene ratio (number of genes /number of genes composing the GO term list). **d**, Gene waves UMAP for highly correlated genes (corr > 0.75), color-coded per type of dynamics (Activated in red, Repressed in green). Bottom left: UMAP of gene waves color-coded per species. Bottom right: cumulative proportions of gene wave dynamics type per species. **e**, Dotplot of DEGs in each indicated comparison of *Mus* time points (columns) per cell class (rows). The size of the dot represents the number of DEGs. **f**, Average wave profile of DEGs identified in E17 vs P1 comparison in *Mus*, for *Mus* (top) and *Acomys* (bottom) for GABAergic neurons. Abbreviations: corr., correlation; DEGs, differentially expressed genes; Gluta, glutamatergic neurons; GABA, GABAergic neurons; IT, intratelencephalically-projecting neurons.

**Extended Data Figure 5.**
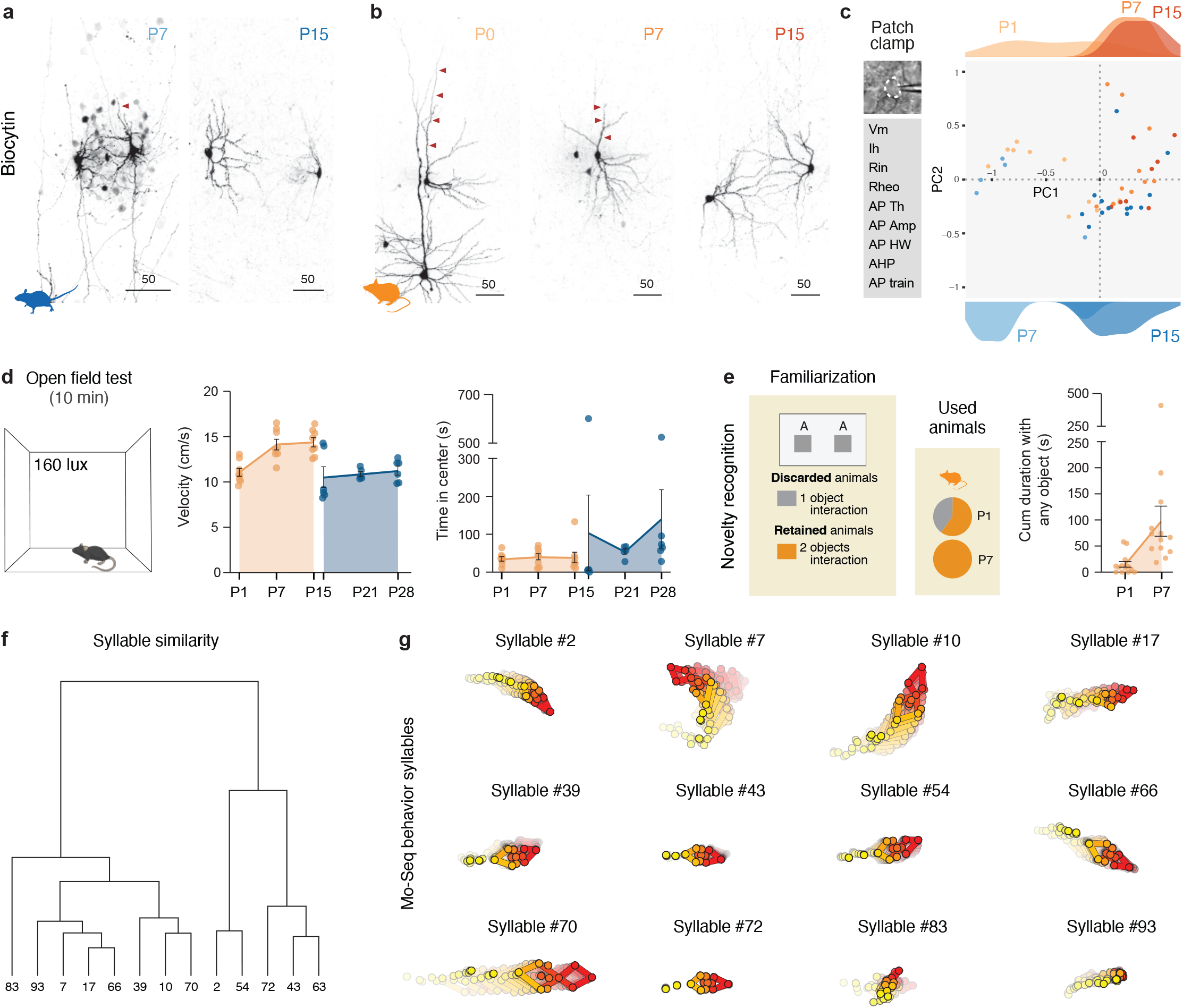
L4 neurons morphology and electrophysiology, and behavioral assessments. **a**, Example images of biocytin-filled L4 neurons in *Mus* at P7 and P15. **b**, Example images of biocytin-filled L4 neurons in *Acomys* at P0, P7 and P15. **c**, PCA analysis of electrophysiological parameters in L4 neurons of *Mus* across time points (blue shades) and *Acomys* across time points (orange shades). Density plots on top and bottom of the PCA plot indicate the density of PC1 values for *Acomys* (top) and *Mus* (bottom) L4 neurons at respective ages (color-coded). **d**, Left, open field test schematics. Middle, Velocity in cm/s of *Acomys* at P1, P7 and P15 and *Mus* at P15, P21 and P28. Right, amount of time spent in the center (in seconds) for *Acomys* at P1, P7 and P15 and *Mus* at P15, P21 and P28. **e**, Left, novelty recognition test schematics. Middle, pie charts summarize the fraction of animals used in the analysis. Only animals that interacted with both objects were considered. Right, plot showing the total time spent interacting with any object for *Acomys* at P1 and P7. **f**, Hierarchical clustering of the syllables showing grouping by similarity. **g**, Average profile of syllables considered for the analysis of MoSeq. Abbreviations: PC, principal component; Vm, resting membrane potential; Ih, hyperpolarization-activated current; Rin, input resistance; Rheo, rheobase; AP Th, action potential threshold; AP amp, action potential amplitude; AP HW, action potential half-width; AHP, hyperpolarization potential; AP train, action potential train.

## Notes

### Competing Interest Statement

The authors have declared no competing interest.

